# Estimating foraging behavior in rodents using a modified paradigm measuring threat imminence dynamics

**DOI:** 10.1101/2023.07.24.550314

**Authors:** Xianzong Meng, Ping Chen, Andor Veltien, Tony Palavra, Sjors In’t Veld, Joanes Grandjean, Judith R Homberg

**Affiliations:** Department of Cognitive Neuroscience, Donders Institute for Brain, Cognition, and Behaviour, Radboud University Medical Centre, 6525 AJ Nijmegen, The Netherlands; Department of Psychiatry, Radboud University Medical Centre, Nijmegen, the Netherlands; Department of Medical Imaging, Radboud University Medical Centre, 6525 GA Nijmegen, The Netherlands

**Keywords:** Neuroscience, Behavioral Neuroscience, Methodologies, Approach-avoid Conflicts

## Abstract

Animals display defensive behaviors in response to threats to avoid danger and approach rewards. In nature, these responses did not evolve alone but are always accompanied by motivational conflict. A semi-naturalistic threat imminence continuum model models the approach-avoidance conflict and is able to integrate multiple defensive behaviors into a single paradigm. However, its comprehensive application is hampered by the lack of a detailed protocol and data about some fundamental factors including sex, age, and motivational level. Here, we modified a previously established paradigm measuring threat imminence continuum dynamics, involving modifications of training and testing protocols, and utilization of commercial materials combined with open science codes, making it more standardized and easier to replicate. We demonstrate that foraging behavior is modulated by age, hunger level, and sex. This paradigm can be used to study defensive behaviors in animals in a more naturalistic manner with relevance to human approach-avoid conflicts and associated psychopathologies.

**Highlights:**

- We provide detailed guidance for setting up a modified paradigm for the threat imminence continuum model with commercial materials and open-source codes.
- We propose a modified paradigm for the threat imminence continuum model to quantify foraging behaviors.
- Our method enables comparison between groups as a function of multiple factors including age and food restriction.

**Graphic Summary:** 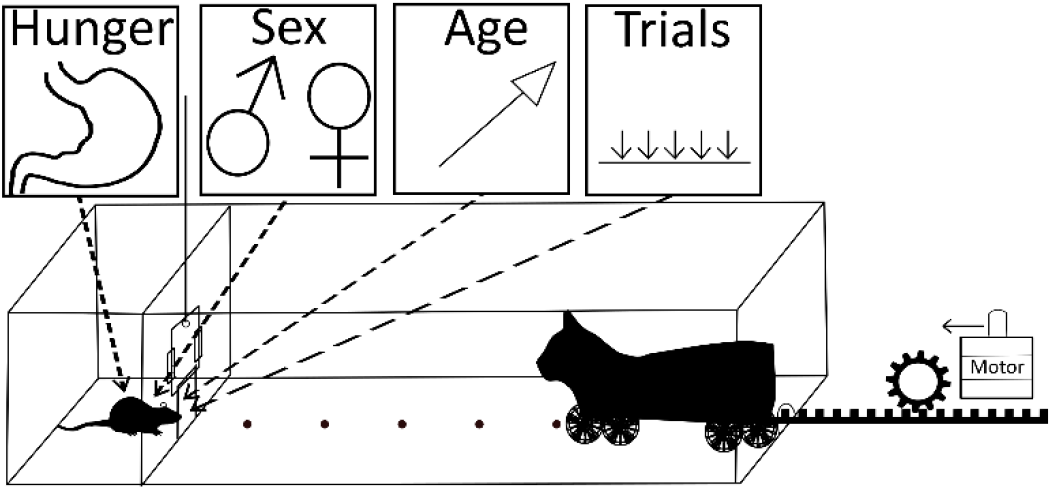

## Introduction

Defensive behaviors strengthen survival and enhance animals’ reproductive rate or capacity (1). Pleasure has been proposed to be evolution’s boldest trick allowing animals and humans to seek primary rewards ensuring survival (2). Importantly, the drive to seek and approach reward did not evolve in isolation but co-evolved with adaptive defensive behaviors such as avoidance to prevent pain/punishment when threats are encountered. When encountering threats and rewards simultaneously, animals are compelled to choose between approach and avoidance behavioral responses (3-6). Maladaptive approach-avoid conflicts have been implicated in the pathophysiology of major depressive disorder (7), and are being used for the measurement of anxiety-related behaviors (8, 9). Proper measurement of the approach-avoidance conflict as it occurs in real life is important for a fundamental understanding of survival behaviors, ecology, as well as human psychopathologies such as anxiety and depression (10, 11).

Unfortunately, current animal paradigms cannot fully satisfy the measurement of the approach-avoidance conflict. Firstly, sometimes the conflict was measured separately (12). For example, in the fear conditioning test (13), robust avoidance responses are triggered but we cannot assess the approach behaviors elicited by rewards. Conditioned suppression of lever pressing for food has been used for Pavlovian fear study including fear incubation (14), fear learning and memory (15). However, conditioned suppression of lever pressing is effective in capturing freeze behavior (15), but cannot encompass the full range of defensive behaviors, e.g. fight. Therefore, our paradigm could help to overcome this limitation. Secondly, the difference between innate fear and learned fear was largely neglected (16, 17). Up to date, despite using different paradigms (e.g., Vogel, Geller-Seifter), most animal paradigms designed for approach-avoidance conflict studies are based on learned fear being associated with both reward and punishment. More importantly, these paradigms failed to address defensive behaviors from a functional aspect. Animals in the wild have the opportunity to display every possible defensive behavior like freezing, flight and even fighting. These behaviors can also influence each other (18). This is not reflected in most current behavioral paradigms for rodents as the animals only get very limited defensive behavioral choices. There is thus a clear need to provide alternative tests to examine behavior in animals across a range of defensive dimensions with a strong translational value for associated studies in humans. Recently, several research groups have addressed this issue by devising novel approach-avoidance conflict tests to assess defensive behavior choices (19-22), the assigned task is foraging, and the threat is induced through either food shock (20-22) or a predator’s odor (19).

To better model approach-avoidance conflict as animals would in the wild, we established a modified threat imminence continuum rat paradigm based on the “threat imminence continuum” theory (23, 24) This theory postulates that defensive behaviors escalate along with three phases, spanning from pre-encounter phase to circa-strike phase: during the *pre-encounter phase,* the threat still has not been detected yet, however, when the likelihood of predation increases or a higher predatory imminence animal perceived, the exploration and foraging behaviors will be reduced (23); during *post-encounter phase*, the threat is perceived but remains remote, which often elicits vigilance and avoidance of the predator; during *circa-strike*, animals will have direct contact with the threat, which elicits an outburst of threat avoidance or combat. To achieve this goal, we establish our paradigm based on the previously published semi-naturalistic Robogator paradigm (4). The paradigm established by Dr. Jeansok Kim’s lab has been used as a reliable method for fear conditioning (25), the integration of fearful information (26), sex effect on predator stimuli (27), and the neurophysiology of predator-induced fear (28). Here, we made some modifications to facilize its replication.

The paradigm consists of a safe nesting area and a foraging area. In the latter, food pellets are made available at various intervals. The threat is represented by a cat-shaped robot resting at the opposite end of the nesting area, propelling toward the rats according to pre-defined rules. By varying the distance the rat needs to navigate to reach the pellet, we can modify the level of threat. This paradigm enables us to examine animal behavior under a more naturalistic condition while retaining the potential for standardization afforded by laboratory experiments. Afterward, to allow a more reproducible implementation of this paradigm across laboratories, reproducible schematics are also provided.

Several modifications were made compared to the original paper we based our study on. Instead of a vague “mild restriction”, our food restriction regimen for the training process was meticulously calculated based on the rats’ actual food consumption under free access conditions. Additionally, we modified the baseline days training from a variable range of 5-7 days to a fixed duration of 6 days. For the behavioral testing, instead of initially placing the food pellet in the middle and subsequently moving its location based on the rats’ performance, we placed the food pellets in order from the nearest position to the nesting area to the farthest. Through this method, we could make the comparison of rats’ performance at different positions, overcoming the limitation of solely focusing on the maximum distance rats could reach. As a result, we could detect more subtle behavioral changes. For instance, some manipulation may not change the maximum distance rats could reach, but can significantly alter the time required for rats to finish foraging at various positions as well as the defensive behavior they exhibit. This contributes to a broader range of experimental possibilities. For statistics, we converted the time data from seconds into rank values, which improved the data’s normality and facilitated its fit into a linear model.

Finally, we set out to investigate the behavioral consequences of important parameters for the optimization of the paradigm, namely sex, age, strain, and motivational levels. We find that sex, age and motivational levels affect rats’ foraging behaviors, and this paradigm can be applied to different rat strains.

## Method

### Preregistration, data and code availability

The statistical method is pre-registered (https://osf.io/v9ypz). The raw data are available under the term of the CC-BY license (https://doi.org/10.34973/38ba-f319, temporary reviewer access link: https://data.donders.ru.nl/login/reviewer-242207846/ZnQ2dnP1fgcjoUxVyvnv0TwQHnRNY0bJMcVnlOol1ec). The code and the processed data to reproduce the assay and the analysis are provided under the terms of the Apache-2.0 license (https://gitlab.socsci.ru.nl/preclinical-neuroimaging/robotmod).

### Subjects

All procedures were carried out in agreement with the current National Research Council Guide for the Care and Use of Laboratory Animals and were approved by the central commission for animal experiments (CCD) in The Hague (the Netherlands), license number AVD10300 2020 9185. All efforts were made to reduce the number of animals used and their suffering. We used mixed-sex groups with a 1:1 sex ratio. Specifically, 12 male and 12 female Sprague Dawley (SD) rats, and 12 male and 12 female Long-Evans (LE) rats were purchased from Charles River (Calco, Italy). For SD rats, two batches of animals were used to test the effect of age on the approach-avoidance conflict in the paradigm. We used 12 postnatal day (PND) 70 rats initially weighing 420–500 g (6 males) and 270-320 g (6 females) and 12 PND 140 rats initially weighing 560-660 g (6 males) and 300-350 g (6 females). For LE rats, we further tested the effect of food restriction on the approach-avoidance conflict. We used 24 PND70 rats initially weighing 360–450 g (12 males) and 220-270 g (12 females). Upon arrival, animals were given two weeks for acclimation to the reversed day-light cycle with free access to food and water. The body weight after this acclimation is considered as initial body weight. Animals were housed in groups of 2 in Macrolon type III cages (42 x 26 x 15 cm) under a 12 h / 12 h reversed day/night cycle (lights off at 8:00 AM) in a temperature-controlled room (22±2°C). Experiments were performed during the dark phase. To motivate the animals for the test, they were subjected to a food restriction regime with *ad libitum* water to maintain 85-95% of their normal weight.

### Threat imminence continuum model setup

The apparatus was assembled by Plexiglas (methyl methacrylate) plates and enhanced by L-shaped anchor plates from the outside with industrial glue (Supplementary Fig 1A). Afterward, another plate serving as a separating plate was positioned inside to create a nesting area (29.21 cm length × 57.12 cm width × 59.69 cm height) and a foraging area (201.93 cm length × 58.42 cm width × 60.96 cm height). On the separating plate, a 10 cm × 10 cm area was laser-cutted at the center of the bottom part, and a remotely controlled door was built to open or close the access from nesting area to foraging area (Supplementary Fig 1B). The robot was attached to a gear rack and the movement was driven by a NEMA17 step motor controlled by an Arduino system. A Raspberry Pi Night Vision Camera (Product code WS-10299) was placed above the center of the cage to record the video (30 frames/s) from both the nesting and foraging areas. All the behavioral tests were performed in a dark experimental room with red light. A customized LEGO-based Robot (66.04 cm length, 17.78 cm width, 15.24 cm height) as shown in Supplementary Fig 1C - 1E, was coded to surge 23 cm (at a velocity of ∼ 75 cm/s) and return to its starting position via an Arduino system (Supplementary Fig 2).

### Food Restriction Regime

Food consumption was measured under a free feeding condition for a week. Afterward, animals received less food based on the calculation from free-feeding conditions throughout the whole experiment. For instance, 90% food restriction means animals could only get 90% of the food they consumed under free access. That is, if the rats could eat 40g food per day under free access conditions, they received 36g food per day to achieve 90% food restriction. During behavioral procedures, food was provided after training or testing. Rats were maintained at 85 - 95% of their body weight.

### Behavioral Procedures

Rats underwent successive stages of habituation days, baseline days, and robot encounter days. The procedure is demonstrated in Supplementary Fig 3.

#### Habituation days

Animals were placed in the nesting area for 30 min/d for 3 consecutive days with three food pellets (grain-based, 0.8 - 1.3 g) inside to acclimatize to the nesting area, during habituation days. The door was closed and animals cannot enter the foraging area.

#### Baseline days

During baseline days, the Robot was absent. After a minute in the nesting area (no food pellets), the door to the foraging area would be opened, and the animal was allowed to explore and search for a food pellet placed 25.4 cm from the nest area (first trial) for 5 min. As soon as the animal took the pellet back inside the nest, the gateway closed. Once the animal finished consuming the pellet, the second foraging trial (with the pellet placed 50.8 cm from the nest area) and then the third foraging trial (with the pellet placed 76.2 cm from the next area) started in the same manner. Animals underwent 6 consecutive baseline days.

#### Robot encounter day

On the robot testing day, the Robot was placed at the opposite end of the foraging area. The food pellet was placed at 25.4 cm, 50.8 cm, 76.2 cm, 101.6 cm, and 127 cm locations in order. After opening the door, each time the animal approached the vicinity (∼1-3cm) of the pellet, the Robot surged 23 cm toward the pellet and returned to its original position. Animals were permitted 3 min to procure the pellet. If animals could procure the food pellet, the door would be closed when animals returned to the nesting area with a food pellet. Subsequently, the food pellet would be placed in the next position. If animals could not procure the food pellet, the door would close with the animal inside the nesting area after 3 min. Under this condition the food pellet was also placed in the next position.

The latency time required for animals to take the food pellet back to the nesting area, the latency required for the animal to go outside of the nesting area, the latency required for animals to approach to food pellet for the first time, and for each position, the number of animals could procure the pellet successfully served as the dependent variables.

## Statistical Analyses

Statistical analysis was optimized on data robot testing day 1 in SD rats. Analysis was performed using R (Version 4.1.2) using the lme4 (lme4_1.1-27.1), multicomp (multcomp_1.4-18), effect size (effectsize_0.6.0.1), and performance (performance_0.8.0) packages. A linear mixed model was built with sex, trials, sex:trials interaction effect, and, if relevant age or food restriction, as fixed effects, and interaction effect with sex or trials, individual intercepts as random effects using the following model: lmer(measured.variable∼ age or food restriction + sex + trials + sex:trials + age or food restriction:sex + age or food restriction:trials + (1|ratID). As a prerequisite to hypothesis testing, the assumptions of normality of the residuals were tested using the performance package. We applied a rank-score conversion (Supplementary Table 1) to enhance the normality of the residuals and minimize the Akaike information criterion (AIC) and Bayesian information criterion (BIC) scores (comparing raw score vs. rank-score, AIC = 785.2 vs 484.8; BIC = 813.6 vs 518.3).

Interpretations of effect size follow Cohen’s 1998 guidelines (Pellow et al., 1985), namely small effect: η2 > 0.02; medium effect: η2 > 0.13; large effect: η2 > 0.26. Results of effect size were presented as effect size (eta-squared, η2) and the 95% confidence intervals (eta-squared upper confidence interval is set to 1 by default).

## Result

### 1. Younger rats finish foraging more rapidly

The foraging behaviors change across the life span in both human beings and rodents (29-32). Firstly we wanted to test the effects of age on animals’ foraging behaviors in the threat imminence continuum model. Two batches of SD rats at different ages (PND 70 and PND 140, n = 12 each, 6/6 m/f) under 90% food restriction were used. We define the success as rats who could successfully bring the food pellet back to the nesting area within the allocated time. Using this metric, we find that older animals had an substantial lower rate of success than younger animals (e.g. PND70 vs PND140 at position 1: 91.7% vs 66.7%; Fig 1A). Next, we tested the overall time needed to reach the food and return to the nesting area, henceforth referred to as foraging. In SD rats data (n = 24, including both PND 70 and PND 140 group), we found that using raw values in seconds broke assumptions of normality of the residuals due to bounded values or missing data in failed trials. To overcome this, we converted raw values to scores (Supplementary Table 1). After correction, we found that older animals needed a longer time to procure the food pellets relative to younger animals (Fig 1B, age effect: η2 = 0.54, [0.28, 1.00]; trial effect η2 = 0.16, [0.04, 1.00]). The position effect denotes longer times to bring pellets back to the nesting areas as a function of the pellet distance from the nest, and will not be discussed further in the text as this is an expected effect in this paradigm. From this experiment, we conclude that younger rats (PND 70) might be more suitable for testing because they avoid a floor effect caused by insufficient success rate, while providing sufficient margins for either improvements or deteriorations in experimental groups.

**Figure 1.**
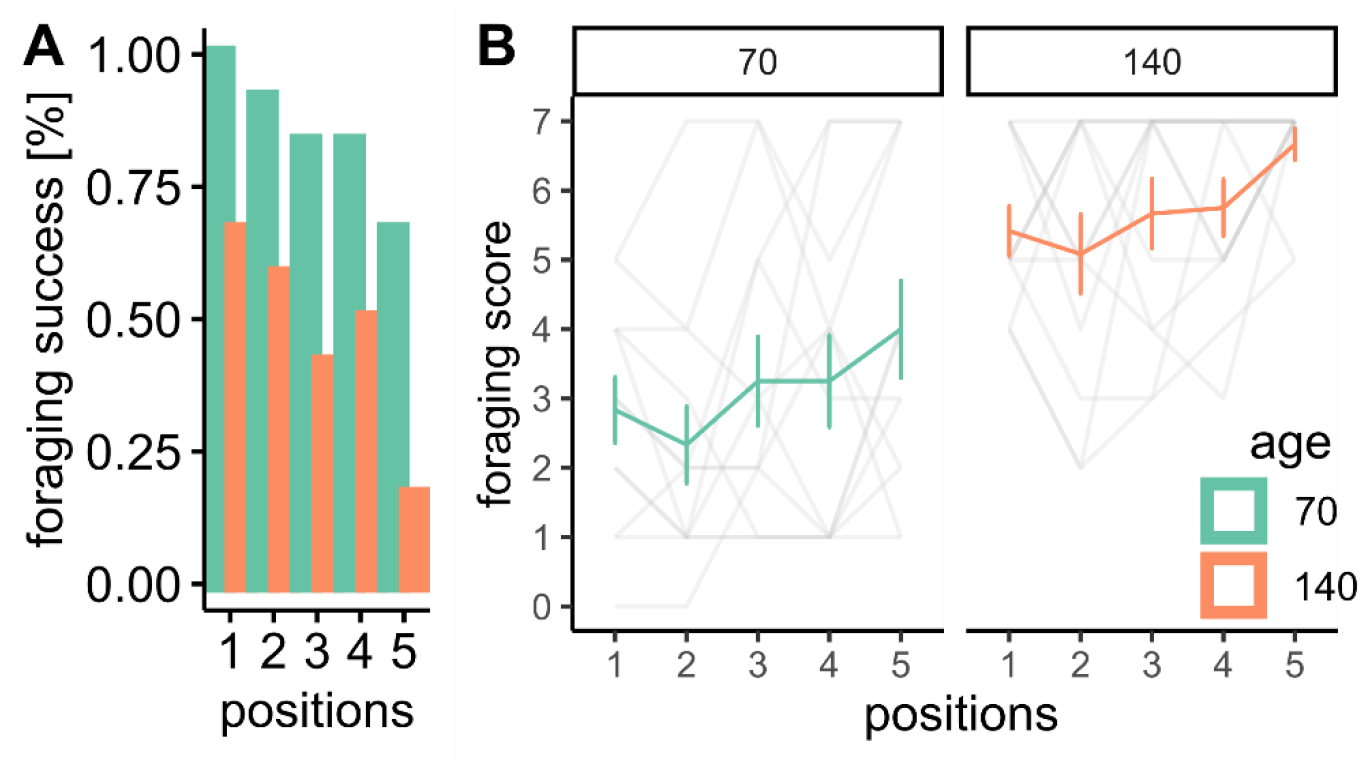
Age effect on rats’ foraging behavior. (A) success rate of younger (PND 70) and older (PND 140) rats in different sex. (B) time to finish foraging. n_PND 70_ = 12 (6/6 (m)male/(f)female), n_PND 140_ = 12 (6/6 m/f). Error bars indicate +/- 1 standard deviation.

### 2. Hunger modulates foraging behavior

In behavioral studies, food restriction is routinely used to initiate or maintain motivational status for animals to engage in different tasks (33), as well as to enhance the motivation to forage for food (34). In our foraging task, a proper food restriction is necessary. However, the extent of food restriction likely influences the foraging behaviors including approach-avoidance decision-making, foraging strategies, speed, hoarding activity, etc. (35-40). Here, we set out to investigate the effect of food restriction. We put LE rats under either 45% (n = 12, male:female ratio = 1:1) or 60% (n = 12, male:female ratio = 1:1) food restriction for 7 days before training and testing the animals with the Robot assay. In this study, we implemented 45% or 60% food restriction regime for rats, which entailed providing rats with either 45% or 60% of the food they consumed under free feeding conditions, thus 45% restriction is associated with a higher hunger level. We find that the success rate increased under the stricter restriction (e.g. 45% restriction vs 60% restriction at position 1 & 5: 100% vs 91.7%; 50% vs 41.7%; Fig 2A). Meanwhile, food restriction has a large effect on foraging performance (restriction effect: η2 = 0.33, [0.07, 1.00]; trial effect: η2 = 0.50, [0.37, 1.00]; sex effect: η2 = 0.33, [0.07, 1.00]), as animals under a 45% restriction regime could take food pellets quicker than animals under 60% restriction regime (Fig 2B).

**Figure 2.**
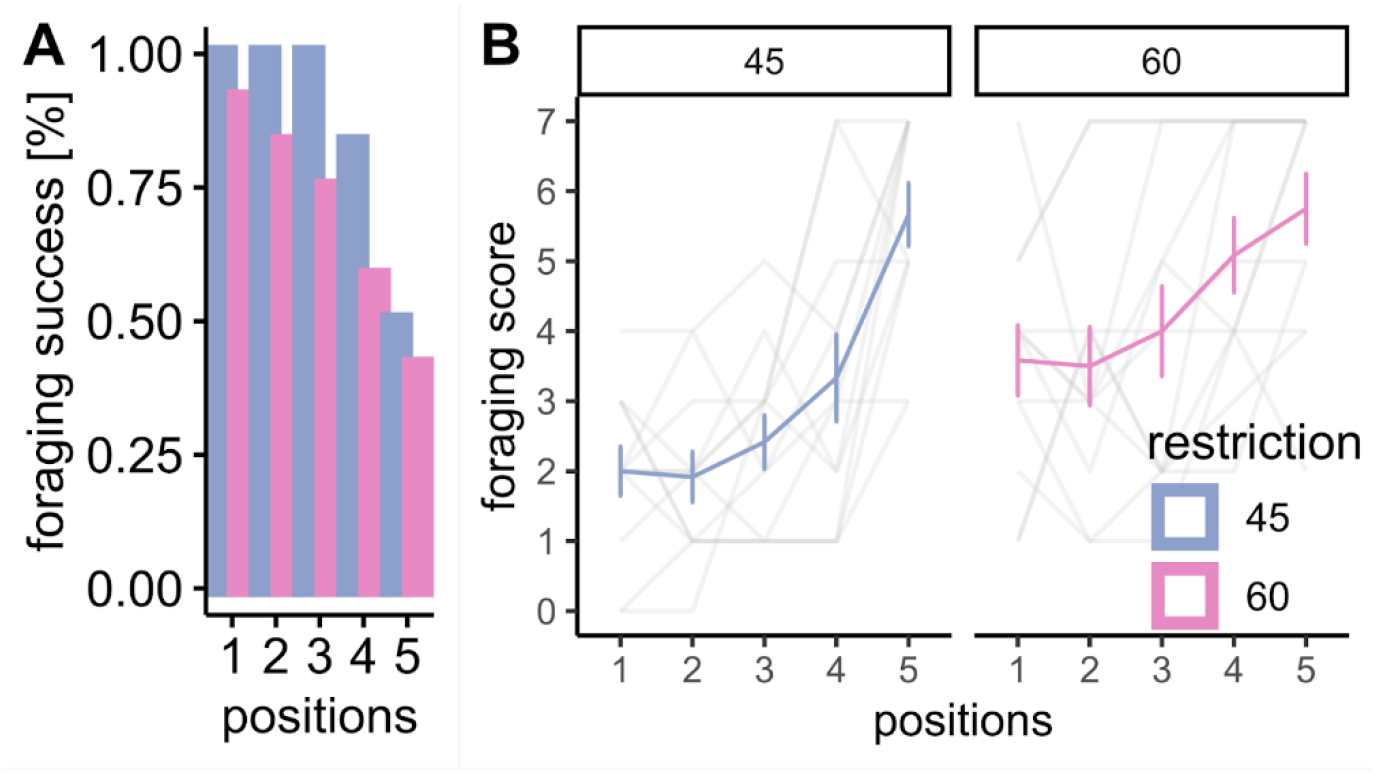
Food restriction effect on rats’ foraging behavior. (A) success rate of foraging under different restrictions in different sexes. (B) time to finish foraging. n_45%_ = 12, n_60%_ = 12, Standard deviation is indicated as the vertical error bars.

### 3. Are female or male rats faster to finish foraging?

As a large sex effect was found from the above food restriction study, we aimed to investigate the effect of sex on test performance. Sex differences in risky-foraging behavior have also been reported when facing aerial predator stimuli (41). However, in our age effect study, only a small sex effect (η2 = 0.03, [0.00, 1.00]) was detected. Meanwhile, the small sex:age interaction effect indicates that age is not a function of sex differences in our paradigm (Fig 3A & 3B).

**Figure 3.**
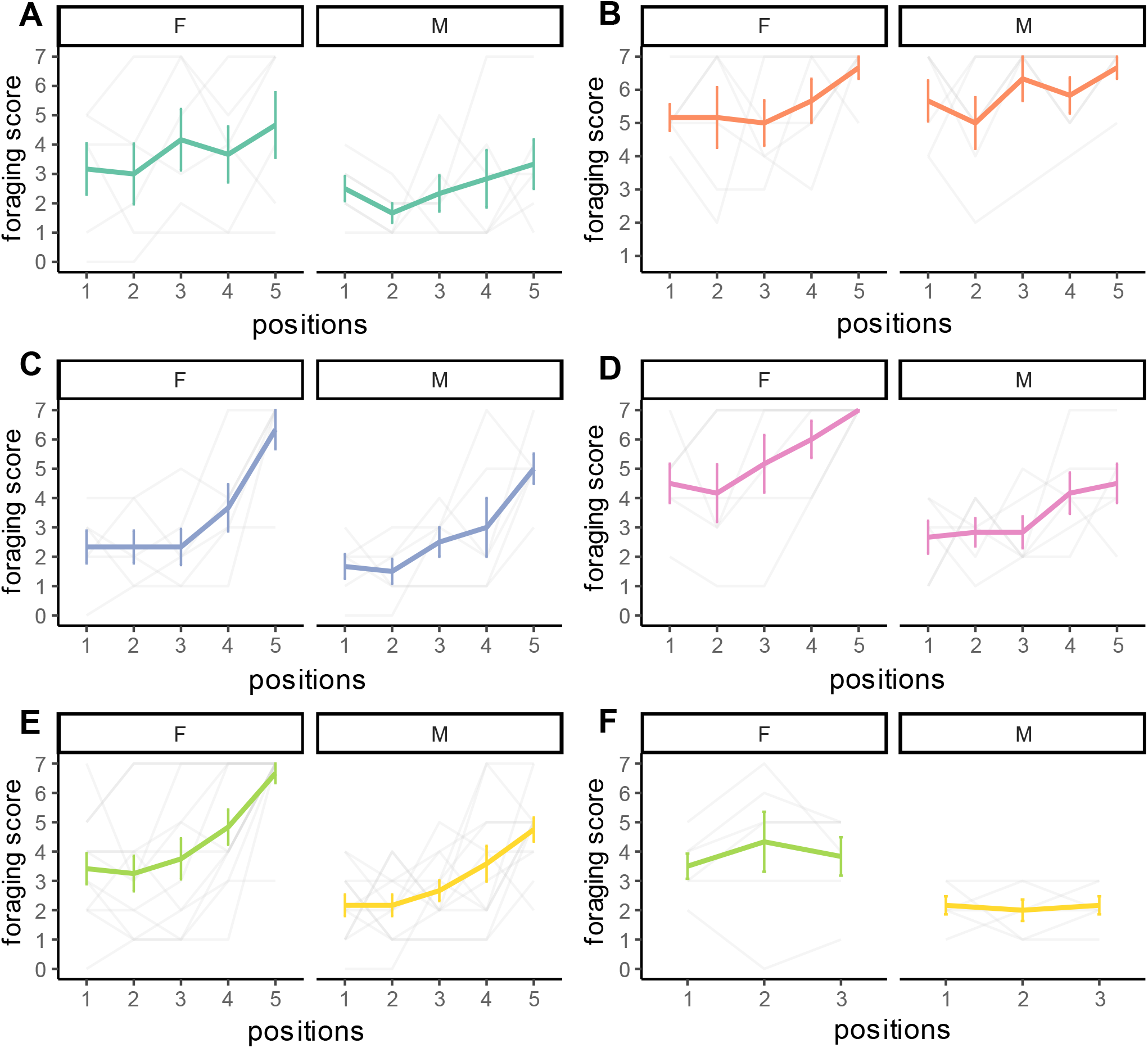
Sex effect on rats’ foraging behavior. (A & B) time to finish foraging at PND70 and PND140, (C & D) time to finish foraging under 45% and 60% food restrictions. (E) general comparison between males and females of time to finish foraging. (F) general comparison between males and females of time to finish foraging at Baseline day 6. n_PND 70_ = 12, n_PND 140_ = 12, n_45%_ = 12, n_60%_ = 12. For Fig 3E, n_male_ = 24 (12/12 45%/60%), n_female_ = 24 (12/12 45%/60%). Standard deviation is indicated as the vertical error bars.

In the food restriction effect study, sex has a larger effect (η2 = 0.33, [0.07, 1.00]) on rats’ foraging performance. At the same time, the small sex:restriction interaction effect suggests that the sex effect is independent from food restrictions. (Fig 3C & 3D). To further investigate the sex effect, we pooled the data from the LE rats together (data from both food restriction regimes), and imported data from Baseline day 6 (the last day of Baseline days). We noticed that female rats needed more time to complete foraging (Fig 3E), even when the robot was not presented (η2 = 0.39, [0.03, 1.00]) (Fig 3F).

### 4. Probing decision-making and risk assessment with leaving and approaching behaviors

The food foraging paradigm plays an important role in behavioral science, and besides foraging behavior itself, the paradigm also involves higher cognitive functions such as decision making and risk assessment. The paradigm we present here allows researchers to measure these advanced behaviors. First, the time the animals need to leave the nesting area (leaving behavior) could reflect animals’ decision making process between starvation risk and predation risk. Secondly, after leaving the nesting area, the time animals need to approach the food pellet for the first time (approaching behavior), could reflect animals’ foraging strategies to prevent or defer the progression of threat.

Meanwhile, these behaviors in foraging could be regulated by animal’s internal state, e.g. satiety state and sex. Therefore we tested the leaving behavior and approaching behavior under different food restriction regimes. For leaving behavior, no obvious deviations were observed (Fig 4A) (food restriction effect η2 = 0.04, [0.00, 1.00], sex effect η2 = 0.04, [0.00, 1.00] and trial effect η2 = 0.07, [0.00, 1.00]). For approaching behavior, a medium sex:restriction effect (η2 = 0.24, [0.02, 1.00]) was observed, indicating that the sex effect and food restriction effect were dependent on each other. Specifically, sex has a medium effect (η2 = 0.18, [0.00, 1.00]) under the 45% restriction condition, and a large effect (η2 = 0.29, [0.00, 1.00]) under the 60% restriction condition (Fig 4B). Meanwhile, restriction has a small effect (η2 = 0.05, [0.00, 1.00]) on males (Fig 4C) but a large effect (η2 = 0.64, [0.26, 1.00]) on females (Fig 4D).

**Figure 4.**
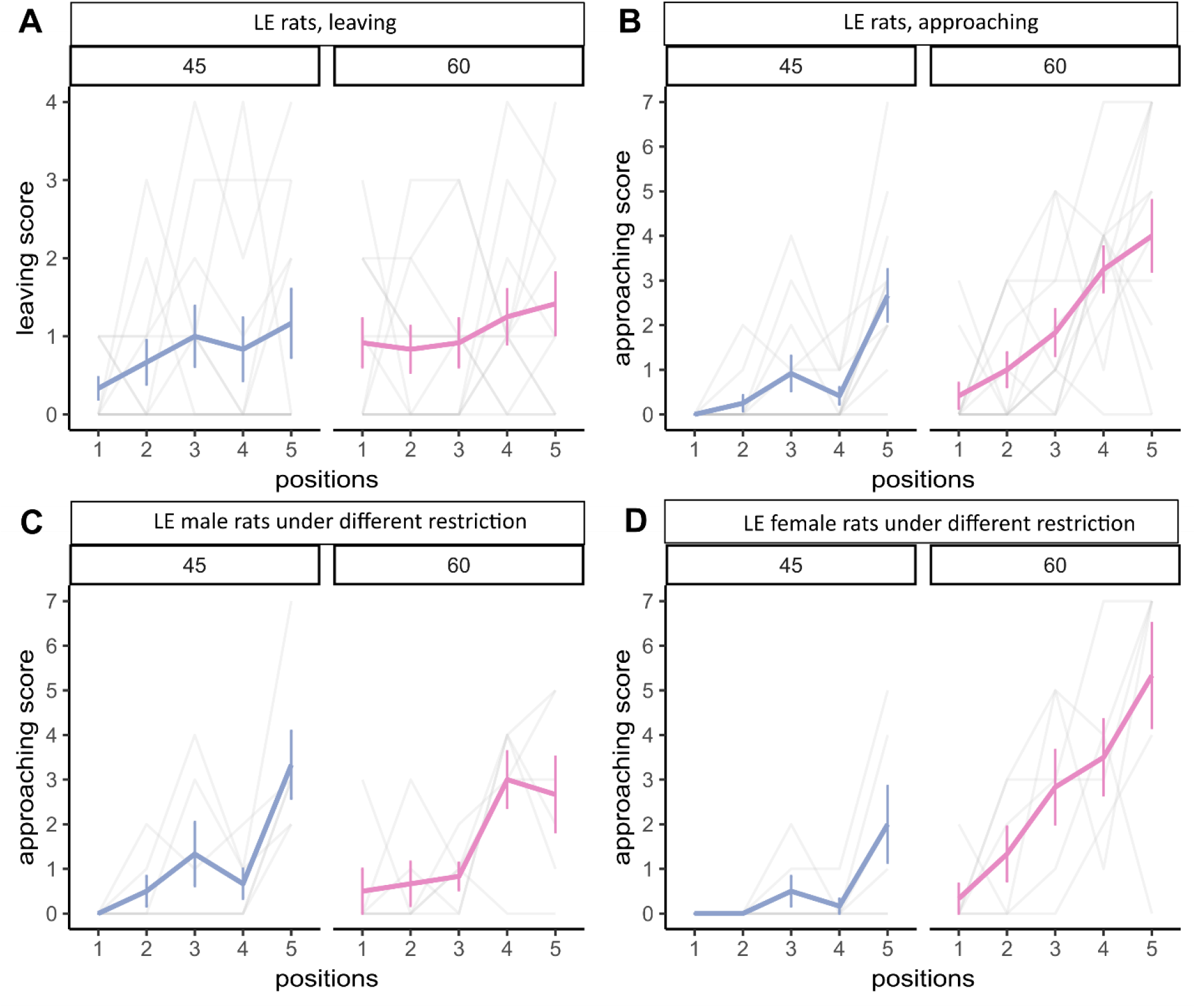
Leaving and approaching behavior during foraging. (A) time to leave the nesting area, (B) time to approach the food pellet for the first time. (C) Approaching score for LE male rats under different food restriction conditions. (D) Approaching score for LE female rats under different food restriction conditions. n_45%_ = 12, n_60%_ = 12. Standard deviation is indicated as the vertical error bars.

## Discussion

Here, we established a modified paradigm based on the threat imminence continuum model to quantify foraging behaviors. Using this paradigm, the effects from some crucial animal parameters including age, sex and food restriction on the foraging behaviors were investigated. Specifically, age and food restriction affected the success rate. We also found that sex influenced performance, however, this seems to be independent of the presence of a threat. We propose that using younger animals with a moderate food restriction regime helps to avoid flooring or ceiling effects, thus allowing the detection of a larger dynamic range of behavior. Finally, to foster further research we provide schematics, code, and assembly instructions to enhance the replication of this modified paradigm.

We found that younger rats could finish foraging faster than older rats. Rats at the age of 2 months are considered as adults because of the maturation of vital systems (42-45). Social maturity starts at the age of 5 and 6 months (42, 43). Although most studies categorized all rats from 2-5 months as adult rats, the decrease of mobility in adulthood cannot be overlooked (32, 46, 47). Meanwhile, clear age-dependent foraging differences have also been reported in rats before, including foraging strategies, food preference and tolerance etc (29, 48). This suggests that the foraging difference we observed was not solely caused by the different locomotion between younger and older rats, but also by a difference in approach-avoidance decision-making. The success rate of younger rats was always higher than older rats in both sexes, indicating that younger rats were more willing to take the risk. However, as to whether the lower success rate in older rats is attributable to reduced motivation for food, changes in punishment sensitivity, or impaired cognitive flexibility still needs to be validated by further experiments.

Food restriction is a common tool used to motivate animals for motivated behaviors of hunger, or learned behaviors (33, 48-52). One alternative method is using palatable rewards (53-56). However, the latter might not generate the level of performance as obtained by food restriction. Palatable food can motivate animals to perform without food deprivation in some circumstances, especially if the task is simple for animals. However, it might take longer for sated animals to learn the task than deprived animals. Sometimes sated animals learn poorly or even refuse to perform the task (51, 53, 57-59). Considering that rats would be foraging in a fearful environment under predation risk, we believe that food restriction could induce a more reliable performance. During the Robot testing process, rats are given 3 min to procure the food pellet from each position. Thus most ideally rats could get food pellets within 3 min after baseline days training. Under 90% food restriction conditions, on baseline day 6 (the last day of baseline training), younger male rats were all able to finish foraging within 3 min. However, all older male rats failed to pass this criterion. Meanwhile, for females, the rates were 66.7% and 16.7% in the younger group and older group, respectively. More importantly, the sex effect in the older group is very small, which is in conflict with reports that female rats are more reluctant to enter a large, novel, open foraging area to procure the food pellets (41, 60-63). We believe that the main reason for this phenomenon is the insufficient food restriction. Because both male and female rats were not motivated enough, many rats refused to go foraging in the risky environment, compromising the sex effect. To improve the sensitivity of our paradigm, we tested two other food restriction regimes, 45% and 60%. We also used a more active rat strain, Long Evans. Data revealed that the success rate gradually reduced along with the distance away from the nesting area as anticipated, and that the more hungry rats were, the quicker they could finish foraging, except when the food pellet was extremely close to the Robot, or in other words, when the threat level was extremely high. It also should be noted that rats under the 45% food restriction could forage very quickly especially from the first four locations in an average of 12 - 35 seconds, which may lead to a ceiling effect. Based on the results above, we recommend using the food restriction at a range around 60%.

As for the sex differences in foraging, when facing the threat, our results indicate that females, in general, take a longer time to finish foraging. This finding is consistent with a previous report that female rats exhibited more potent fear responses to aerial threats (41). However, this phenomenon seems to be threat-independent (even when the Robot is not present). Therefore, we further would like to conclude that our assay may not be a very robust tool to tweak out sex differences because of the fundamental sex differences on rats’ foraging behaviors. Meanwhile, it also has been reported that male and female rats would use different strategies when the threat was integrated into their life. That is, male rats would be more willing to put effort in risky foraging because it can increase mate access (64, 65), whereas female rats would slash their requirement of food in order to avoid aversive stimuli (60). However, whether it is due to the alteration of fear perception or adaptation of strategy still needs to be investigated.

The effect from above mentioned factors on rats’ leaving and approaching behaviors was also investigated, which enables us to explore more cognitive functions, including risky decision making and risk assessment. When the door was opened, animals would be able to see the Robot and were required to make a decision whether or not to leave the nesting area for foraging. As proposed by Fendt et al. (18), the decision-making of animals is based on the integration of various factors like hunger, distance of threat, etc. In the present study, two internal factors, hunger level (45% and 60%) and sex, and one external factor distance of threat was studied. Results indicate that all above mentioned factors only have a small effect on rats’ foraging decision. Interestingly, it has been previously reported that males exhibit greater risk-taking than females in decision making when involving the risk of punishment (64). However, in our paradigm, sex only exhibited a small effect on foraging decision (leaving behavior). Therefore, the possibility cannot be excluded that once the hunger level has reached a certain threshold, the effect from sex and distance of threat on the decision-making process will have a very limited impact. Thereby, optimizing the food restriction regime is highly recommended when using this paradigm for decision-making studies. For approach behavior, as reported by Kim et al in their paper using a similar paradigm (4), with the appearance of Robot, rats do not simply give up foraging, but try to use different strategies and repeated efforts to procure the food pellets. This suggests the involvement of risk assessment. Results revealed a medium sex:restriction interaction effect, and further analysis found that under a milder restriction regime, sex could have a larger effect on the risk assessment process. Furthermore, the results suggest that food restriction has a more profound effect on females compared with males. The mechanism behind this phenomenon still needs deeper investigation. However, it should be noted that ovariectomy in adult female rats did not alter the risky foraging strategies, implying that these changes are prepared during early development (60). Here, we assume that the difference in threat perception, information integration, and ghrelin secretion between males and females might contribute to the changes. Yet, this also still needs to be further investigated.

For specific defensive behaviors, In this paper, we did not directly measure the defensive behaviors, however, our paradigm allows the recording of the animals’ performance during experimental sessions, and through tools like EthoVision or DeepLabCut, the various defensive responses exhibited by the animals, including freezing, flight, and fight, can be quantified and analyzed.

Some limitations should be taken into account when interpreting the data. Firstly, the different visual acuities between males and females might also account for the observed sex differences (66, 67). Secondly, after 6 days of baseline training, the possibility cannot be ruled out that rats decided to leave the nesting area too quickly to clearly see or notice the appearance of the Robot. Therefore, for experiments focusing on decision-making processes, we recommend replacing the separating plate with a transparent one, and triggering the Robot a few times before behavioral testing to allow rats to perceive the threat in advance. Thirdly, in this study, the risk assessment was evaluated by measuring the time it took for rats to attempt to procure the food pellet for the very first time after leaving their nesting area. This method for evaluating risk assessment is simple and direct, however, it should not be overlooked that approaching to food behavior is regulated by some other factors, e.g. social cues (68). Therefore, the specificity of the risk assessment measure might need to be improved, which could be achieved by analyzing specific behaviors of the rats, including head-dipping, rearing, grooming, etc., especially Stretch-attend posture, which is currently the most prevalent method (69-74). Furthermore, in this paradigm, the food will be provided after training or behavioral test, that makes our paradigm an “open economy” type, which narrows the detection range of behaviors (75) and excludes information from temporal aspects of fear and anxiety (76). As proposed by Collier et al., (77) a “closed economy” paradigm in which animals obtain their daily food exclusively from operant process and typically live in the operant chambers could provide be a more holistic approach to for rodent behavior study. By introducing aversive components to the “closed economy”, “risky closed economy” was developed (78, 79), through which the information from a spatiotemporal dimension of fear and anxiety could be obtained. And its naturalistic qualities could facilitate the assessment of fear and anxiety in decision-making (5). However, some practical limitations of “Risky Closed Economy” might hinder its application, e.g. it requires substantial amount of time and space and rats’ social interaction may be reduced (76). Besides, measuring stress hormones like corticosterone could provide a fuller picture of rats’ fear level as well as validate how much fear the robot triggered (80, 81).

In conclusion, we propose a modified paradigm based on threat imminence continuum model which could enable us to quantify foraging behaviors along with our paradigm. Some key factors influencing foraging and approach-avoidance behavior were investigated. Results suggested that age, sex and hunger level could affect rats foraging performance, as well as risk assessment. However, whether these factors could also affect risky decision making still needs to be further investigated. Besides, our paradigm could serve as a powerful tool to exploit the full potential of comprehensive behavioral phenotyping of rats’ foraging under threat in a semi-naturalistic environment.

## Acknowledgements

This project is funded by Radboudumc internal budget.

## Author contributions

J.H. and J.G. design and supervision. X.M., A.V. experimental cage building. X.M., T.P. data collection, X.M., J.G., P.C., data analysis. X.M., S.V. Python coding. J.H., J.G., X.M., P.C., A.V. paper writing and editing. All authors had full access to all of the data in the study and accept responsibility for the decision to submit for publication.

## Supplemental information

**Supplementary Fig 1.**
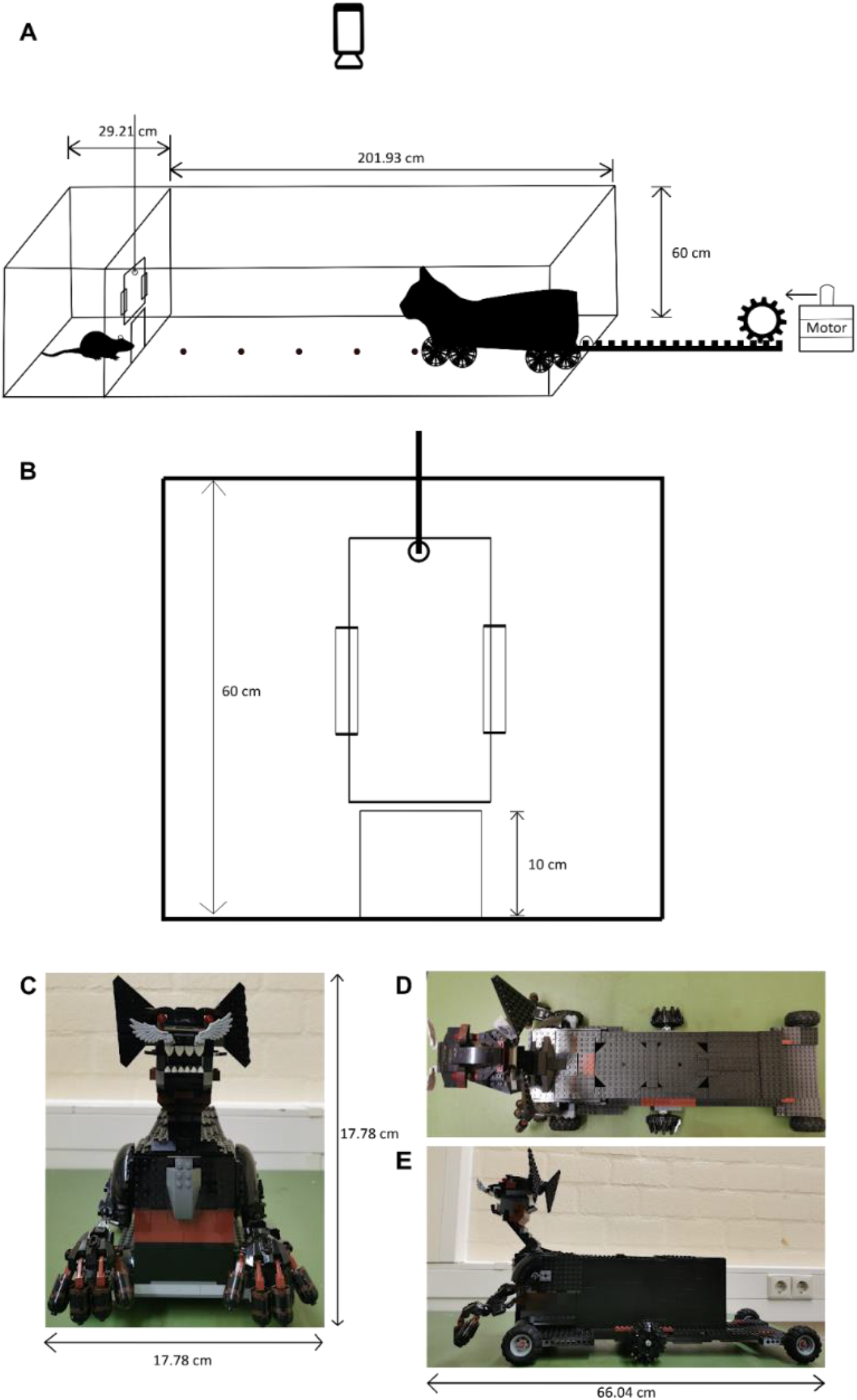
Schematics of the paradigm setup. (A) Overview of the paradigm. (B) Controlled-door on the separating plate. (C) The front view of the Robot. (D) The top view of the Robot. (E) The side view of the Robot. Note: for building the setup, we recommend controlling the door manually to avoid causing injury and extra stress to rats, because rats always poke their head out before going foraging and sometimes leave their tails outside the nesting area when taking food back.

**Supplementary Fig 2.**
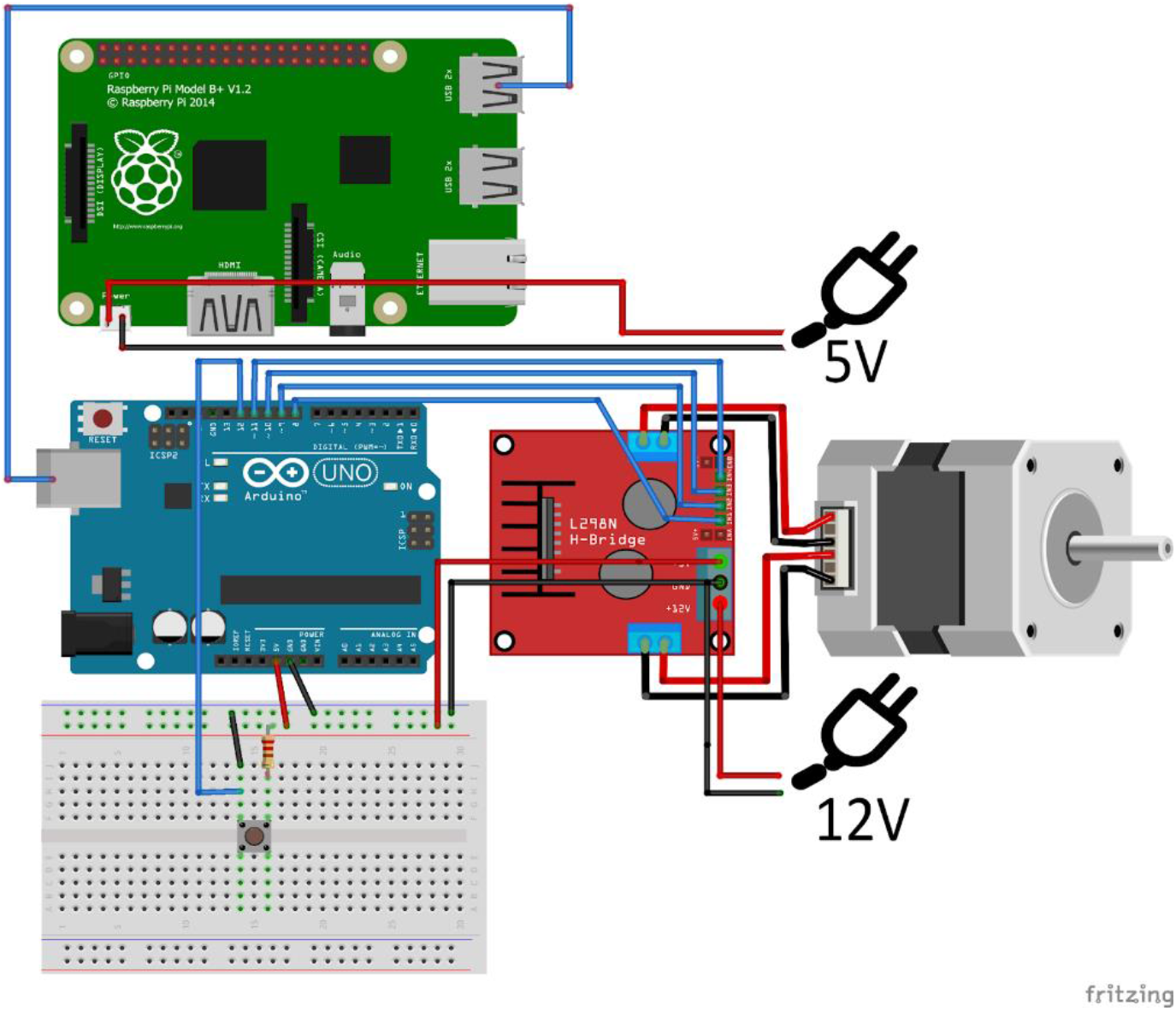
Schematics of connections between the Arduino board, Raspberry Pi and Step motor.

**Supplementary Fig 3.**
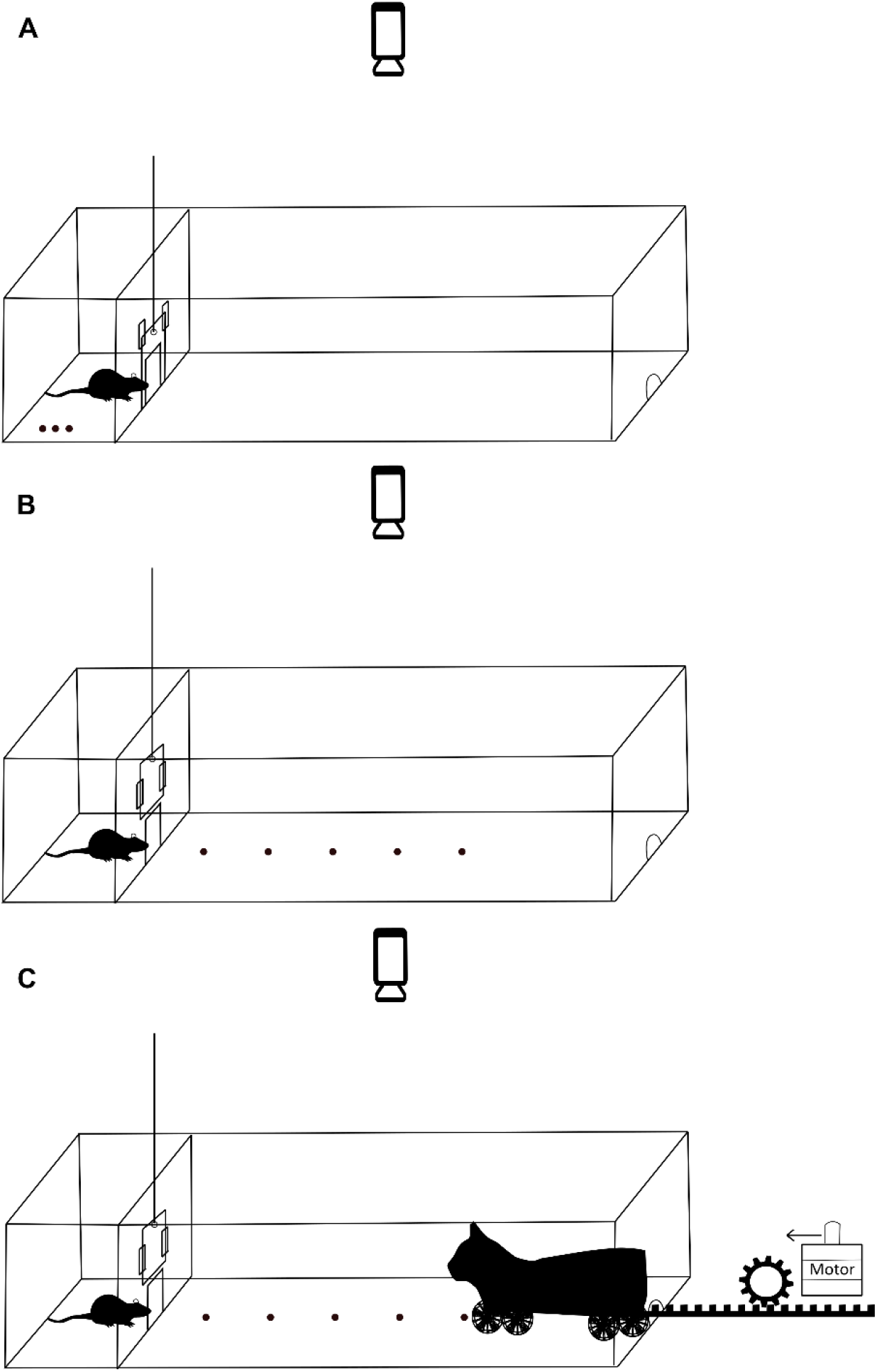
Schematics of behavioral procedures. (A) Habituation. (B) Baseline days. (C) Robot testing day.

**Supplementary Table 1.** Rank-score conversion.

| animals performance | score value |
| --- | --- |
| failed to complete the task | 7 |
| complete the task using more than 200 sec | 6 |
| complete the task within 200 - 100 sec | 5 |
| complete the task within 100 - 50 sec | 4 |
| complete the task within 50 - 25 sec | 3 |
| complete the task within 25 - 12.5 sec | 2 |
| complete the task within 12.5 - 6.25 sec | 1 |
| complete the task within 6.25 - 0 sec | 0 |

**Supplementary videos.**
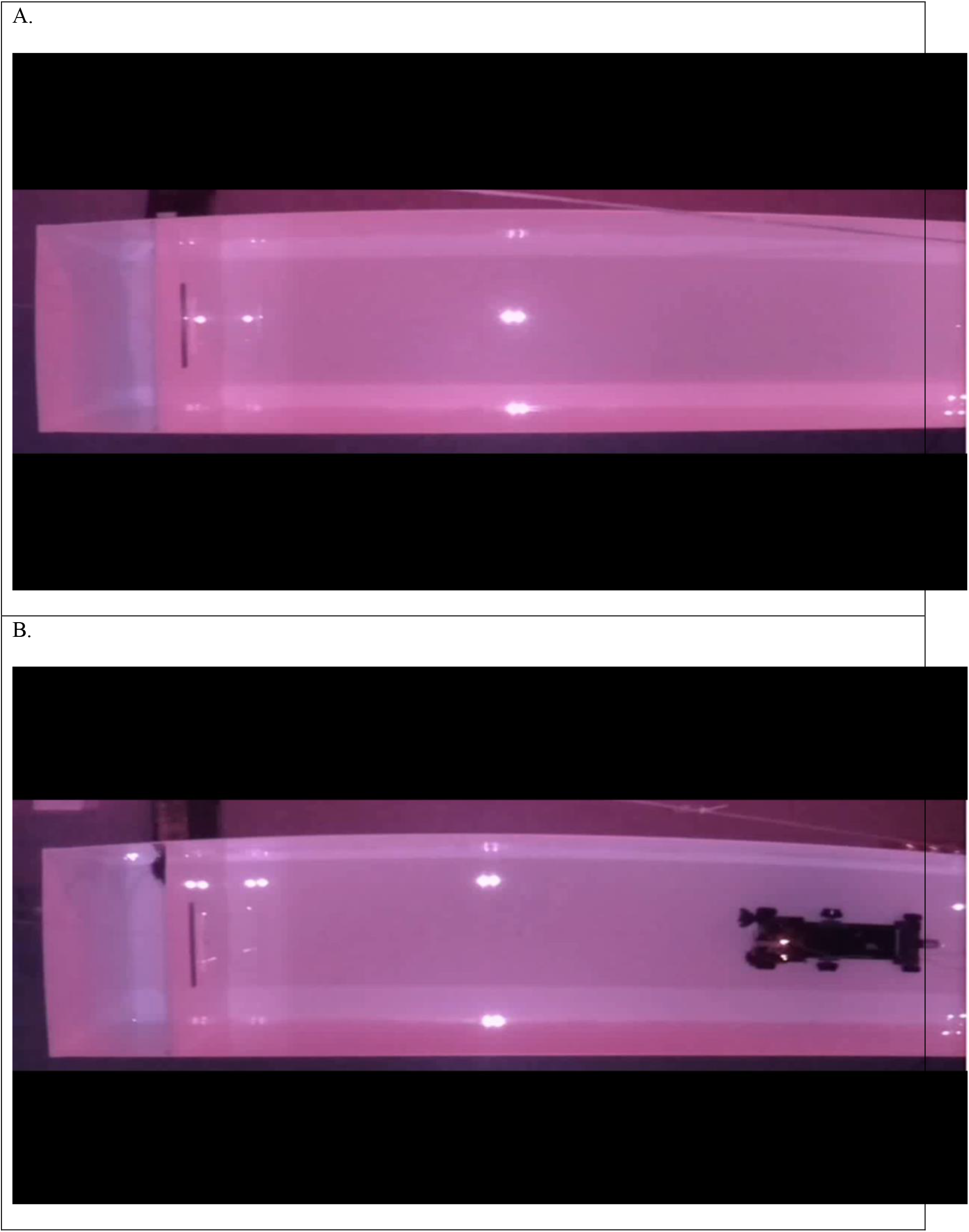
(A). Representative video for baseline days, food pellet was placed at position 3. (B) Representative video for Robot testing days, food pellet was placed at position 3. Note: the video has been speedup for demonstration purpose.

## Reference

1. Williams JL. Historical and Contemporary Approaches to the Study of Defense in Animals. The Psychological Record. 1991;41(2):153–8.

2. Schultz W. Neuronal Reward and Decision Signals: From Theories to Data. Physiological reviews. 2015;95(3):853–951.

3. Amir A, Lee SC, Headley DB, Herzallah MM, Pare D. Amygdala Signaling during Foraging in a Hazardous Environment. The Journal of neuroscience: the official journal of the Society for Neuroscience. 2015;35(38):12994–3005.

4. Choi JS, Kim JJ. Amygdala regulates risk of predation in rats foraging in a dynamic fear environment. Proceedings of the National Academy of Sciences of the United States of America. 2010;107(50):21773–7.

5. Mobbs D, Trimmer PC, Blumstein DT, Dayan P. Foraging for foundations in decision neuroscience: insights from ethology. Nature reviews Neuroscience. 2018;19(7):419–27.

6. Hayden BY, Walton ME. Neuroscience of foraging. Frontiers in neuroscience. 2014;8:81.

7. Ironside M, Amemori KI, McGrath CL, Pedersen ML, Kang MS, Amemori S, et al. Approach-Avoidance Conflict in Major Depressive Disorder: Congruent Neural Findings in Humans and Nonhuman Primates. Biological psychiatry. 2020;87(5):399–408.

8. La-Vu M, Tobias BC, Schuette PJ, Adhikari AJFibn. To approach or avoid: an introductory overview of the study of anxiety using rodent assays. 2020;14:145.

9. Aupperle RL, McDermott TJ, White E, Kirlic N. The neuropsychology of anxiety: An approach– avoidance decision-making framework. 2023.

10. Rusconi F, Rossetti MG, Forastieri C, Tritto V, Bellani M, Battaglioli E. Preclinical and clinical evidence on the approach-avoidance conflict evaluation as an integrative tool for psychopathology. Epidemiology and psychiatric sciences. 2022;31:e90.

11. Kirlic N, Young J, Aupperle RL. Animal to human translational paradigms relevant for approach avoidance conflict decision making. Behaviour research and therapy. 2017;96:14–29.

12. Hu H. Reward and Aversion. Annual review of neuroscience. 2016;39:297–324.

13. Curzon P, Rustay NR, Browman KE. Frontiers in Neuroscience Cued and Contextual Fear Conditioning for Rodents. In: Buccafusco JJ, editor. Methods of Behavior Analysis in Neuroscience. Boca Raton (FL): CRC Press/Taylor & Francis Copyright © 2009, Taylor & Francis Group, LLC.; 2009.

14. Pickens CL, Navarre BM, Nair SG. Incubation of conditioned fear in the conditioned suppression model in rats: role of food-restriction conditions, length of conditioned stimulus, and generality to conditioned freezing. Neuroscience. 2010;169(4):1501–10.

15. McDannald MA, Galarce EM. Measuring Pavlovian fear with conditioned freezing and conditioned suppression reveals different roles for the basolateral amygdala. Brain research. 2011;1374:82–9.

16. Corcoran KA, Quirk GJ. Activity in prelimbic cortex is necessary for the expression of learned, but not innate, fears. The Journal of neuroscience: the official journal of the Society for Neuroscience. 2007;27(4):840–4.

17. Silva BA, Gross CT, Gräff J. The neural circuits of innate fear: detection, integration, action, and memorization. Learning & memory (Cold Spring Harbor, NY). 2016;23(10):544–55.

18. Fendt M, Parsons MH, Apfelbach R, Carthey AJR, Dickman CR, Endres T, et al. Context and trade-offs characterize real-world threat detection systems: A review and comprehensive framework to improve research practice and resolve the translational crisis. Neuroscience and biobehavioral reviews. 2020;115:25–33.

19. Engelke DS, Zhang XO, O’Malley JJ, Fernandez-Leon JA, Li S, Kirouac GJ, et al. A hypothalamic-thalamostriatal circuit that controls approach-avoidance conflict in rats. Nature communications. 2021;12(1):2517.

20. Illescas-Huerta E, Ramirez-Lugo L, Sierra RO, Quillfeldt JA, Sotres-Bayon F. Conflict Test Battery for Studying the Act of Facing Threats in Pursuit of Rewards. Frontiers in neuroscience. 2021;15:645769.

21. Fernandez-Leon JA, Engelke DS, Aquino-Miranda G, Goodson A, Rasheed MN, Do Monte FH. Neural correlates and determinants of approach-avoidance conflict in the prelimbic prefrontal cortex. eLife. 2021;10.

22. Bravo-Rivera H, Rubio Arzola P, Caban-Murillo A, Vélez-Avilés AN, Ayala-Rosario SN, Quirk GJ. Characterizing Different Strategies for Resolving Approach-Avoidance Conflict. Frontiers in neuroscience. 2021;15:608922.

23. Fanselow MS, Lester LS. A Functional Behavioristic Approach to Aversively Motivated Behavior:: Predatory Imminence as a Determinant of the Topography of Defensive Behavior. Evolution and learning: Psychology Press; 2013. p. 185-212.

24. Bouton ME, Mineka S, Barlow DHJPr. A modern learning theory perspective on the etiology of panic disorder. 2001;108(1):4.

25. Zambetti PR, Schuessler BP, Lecamp BE, Shin A, Kim EJ, Kim JJ. Ecological analysis of Pavlovian fear conditioning in rats. Communications biology. 2022;5(1):830.

26. Kong MS, Kim EJ, Park S, Zweifel LS, Huh Y, Cho J, et al. ‘Fearful-place’ coding in the amygdala-hippocampal network. eLife. 2021;10.

27. Zambetti PR, Schuessler BP, Kim JJ. Sex Differences in Foraging Rats to Naturalistic Aerial Predator Stimuli. iScience. 2019;16:442–52.

28. Kim EJ, Kong MS, Park SG, Mizumori SJY, Cho J, Kim JJ. Dynamic coding of predatory information between the prelimbic cortex and lateral amygdala in foraging rats. Science advances. 2018;4(4):eaar7328.

29. Clark DAJTJoAE. Age-and sex-dependent foraging strategies of a small mammalian omnivore. 1980:549-63.

30. Mata R, Wilke A, Czienskowski UJFin. Foraging across the life span: is there a reduction in exploration with aging? 2013;7:53.

31. Shoji H, Miyakawa TJNr. Age-related behavioral changes from young to old age in male mice of a C57 BL/6J strain maintained under a genetic stability program. 2019;39(2):100-18.

32. Boguszewski P, Zagrodzka JJBbr. Emotional changes related to age in rats—a behavioral analysis. 2002;133(2):323–32.

33. Dietze S, Lees KR, Fink H, Brosda J, Voigt J-PJA. Food deprivation, body weight loss and anxiety-related behavior in rats. 2016;6(1):4.

34. Selakovic D, Joksimovic JJSJoE, Research C. Behavioural effects of short-term total food restriction in rats. 2014;15(3):129–37.

35. Hernández MC, Navarro-Castilla Á, Barja IJPo. Wood mouse feeding effort and decision-making when encountering a restricted unknown food source. 2019;14(6):e0212716.

36. Gutman R, Yosha D, Choshniak I, Kronfeld-Schor NJP, behavior. Two strategies for coping with food shortage in desert golden spiny mice. 2007;90(1):95–102.

37. Heinz DE, Schöttle VA, Nemcova P, Binder FP, Ebert T, Domschke K, et al. Exploratory drive, fear, and anxiety are dissociable and independent components in foraging mice. 2021;11(1):318.

38. Burnett CJ, Li C, Webber E, Tsaousidou E, Xue SY, Brüning JC, et al. Hunger-driven motivational state competition. 2016;92(1):187–201.

39. Dailey MJ, Bartness TJJBr. Arcuate nucleus destruction does not block food deprivation-induced increases in food foraging and hoarding. 2010;1323:94–108.

40. Keen-Rhinehart E, Dailey MJ, Bartness TJPTotRSBBS. Physiological mechanisms for food-hoarding motivation in animals. 2010;365(1542):961-75.

41. Zambetti PR, Schuessler BP, Kim JJJI. Sex differences in foraging rats to naturalistic aerial predator stimuli. 2019;16:442–52.

42. Sengupta PJIjopm. The laboratory rat: relating its age with human’s. 2013;4(6):624.

43. Andreollo NA, Santos EFd, Araújo MR, Lopes LRJAABdCD. Rat’s age versus human’s age: what is the relationship? 2012;25:49–51.

44. Quinn RJN. Comparing rat’s to human’s age: how old is my rat in people years? 2005;21(6):775.

45. Klein ZA, Romeo RDJH, behavior. Changes in hypothalamic–pituitary–adrenal stress responsiveness before and after puberty in rats. 2013;64(2):357–63.

46. Sudakov SK, Alekseeva EV, Nazarova GA, Bashkatova VGJA. Age-related individual behavioural characteristics of adult wistar rats. 2021;11(8):2282.

47. Arrant AE, Schramm-Sapyta NL, Kuhn CMJBbr. Use of the light/dark test for anxiety in adult and adolescent male rats. 2013;256:119–27.

48. Bert B, Harms S, Langen B, Fink HJJovp, therapeutics. Clomipramine and selegiline: do they influence impulse control? 2006;29(1):41–7.

49. Carlini V, Martini A, Schiöth H, Ruiz R, De Cuneo MF, De Barioglio SJN. Decreased memory for novel object recognition in chronically food-restricted mice is reversed by acute ghrelin administration. 2008;153(4):929–34.

50. Conrad CDJPiN-P, Psychiatry B. A critical review of chronic stress effects on spatial learning and memory. 2010;34(5):742–55.

51. Kant GJ, Yen MH, D’Angelo PC, Brown AJ, Eggleston TJPB, Behavior. Maze performance: a direct comparison of food vs. water mazes. 1988;31(2):487–91.

52. Heiderstadt K, McLaughlin R, Wrighe D, Walker S, Gomez-Sanchez CJLa. The effect of chronic food and water restriction on open-field behaviour and serum corticosterone levels in rats. 2000;34(1):20-8.

53. Toth LA, Gardiner TWJJotAAfLAS. Food and water restriction protocols: physiological and behavioral considerations. 2000;39(6):9–17.

54. Young PT, Shuford Jr EJJoC, Psychology P. Quantitative control of motivation through sucrose solutions of different concentrations. 1955;48(2):114.

55. Carroll ME, Lac ST, Nygaard SLJP. A concurrently available nondrug reinforcer prevents the acquisition or decreases the maintenance of cocaine-reinforced behavior. 1989;97:23–9.

56. Ray SJAT. An alternative to water deprivation techniques in animal learning studies. 1998.

57. Hughes JE, Amyx H, Howard JL, Nanry KP, Pollard GTJLas. Health effects of water restriction to motivate lever-pressing in rats. 1994;44(2):135–40.

58. Collier G, Levitsky DJJoc, psychology p. Defense of water balance in rats: behavioral and physiological responses to depletion. 1967;64(1):59.

59. Lattal KA, Williams AMJJotEaob. Body weight and response acquisition with delayed reinforcement. 1997;67(1):131–43.

60. Pellman BA, Schuessler BP, Tellakat M, Kim JJJe. Sexually dimorphic risk mitigation strategies in rats. 2017;4(1).

61. Carrier N, Kabbaj MJN. Sex differences in the antidepressant-like effects of ketamine. 2013;70:27–34.

62. Jolles JW, Boogert NJ, van den Bos RJRSOS. Sex differences in risk-taking and associative learning in rats. 2015;2(11):150485.

63. Turner KM, Burne THJPo. Comprehensive behavioural analysis of Long Evans and Sprague-Dawley rats reveals differential effects of housing conditions on tests relevant to neuropsychiatric disorders. 2014;9(3):e93411.

64. Ellis BJ, Figueredo AJ, Brumbach BH, Schlomer GLJHN. Fundamental dimensions of environmental risk: The impact of harsh versus unpredictable environments on the evolution and development of life history strategies. 2009;20:204–68.

65. West-Eberhard MJ. Darwin’s forgotten idea: the social essence of sexual selection. Neuroscience and biobehavioral reviews. 2014;46 Pt 4:501–8.

66. Seymoure P, Juraska JM. Vernier and grating acuity in adult hooded rats: the influence of sex. Behavioral neuroscience. 1997;111(4):792–800.

67. Juraska JM. Changes in sex differences in neuroanatomical structure and cognitive behavior across the life span. Learning & memory (Cold Spring Harbor, NY). 2022;29(9):340–8.

68. Hall FS, Humby T, Wilkinson LS, Robbins TW. The effects of isolation-rearing of rats on behavioural responses to food and environmental novelty. Physiology & behavior. 1997;62(2):281–90.

69. Holly KS, Orndorff CO, Murray TA. MATSAP: An automated analysis of stretch-attend posture in rodent behavioral experiments. Scientific reports. 2016;6:31286.

70. Albrechet-Souza L, Cristina de Carvalho M, Rodrigues Franci C, Brandão ML. Increases in plasma corticosterone and stretched-attend postures in rats naive and previously exposed to the elevated plus-maze are sensitive to the anxiolytic-like effects of midazolam. Hormones and behavior. 2007;52(2):267–73.

71. Magara S, Holst S, Lundberg S, Roman E, Lindskog M. Altered explorative strategies and reactive coping style in the FSL rat model of depression. Frontiers in behavioral neuroscience. 2015;9:89.

72. Coimbra NC, Paschoalin-Maurin T, Bassi GS, Kanashiro A, Biagioni AF, Felippotti TT, et al. Critical neuropsychobiological analysis of panic attack- and anticipatory anxiety-like behaviors in rodents confronted with snakes in polygonal arenas and complex labyrinths: a comparison to the elevated plus- and T-maze behavioral tests. Revista brasileira de psiquiatria (Sao Paulo, Brazil: 1999). 2017;39(1):72-83.

73. Rodgers RJ, Haller J, Holmes A, Halasz J, Walton TJ, Brain PF. Corticosterone response to the plus-maze: high correlation with risk assessment in rats and mice. Physiology & behavior. 1999;68(1-2):47–53.

74. Reis FM, Albrechet-Souza L, Franci CR, Brandão ML. Risk assessment behaviors associated with corticosterone trigger the defense reaction to social isolation in rats: role of the anterior cingulate cortex. Stress (Amsterdam, Netherlands). 2012;15(3):318–28.

75. Pellman BA, Kim JJJTin. What can ethobehavioral studies tell us about the brain’s fear system? 2016;39(6):420–31.

76. Schuessler BP, Zambetti PR, Kukuoka KM, Kim EJ, Kim JJ. The Risky Closed Economy: A Holistic, Longitudinal Approach to Studying Fear and Anxiety in Rodents. Frontiers in behavioral neuroscience. 2020;14:594568.

77. Collier G, Hirsch E, Hamlin PHJP, Behavior. The ecological determinants of reinforcement in the rat. 1972;9(5):705–16.

78. Fanselow MS, Lester LS, Helmstetter FJJJoteaob. Changes in feeding and foraging patterns as an antipredator defensive strategy: a laboratory simulation using aversive stimulation in a closed economy. 1988;50(3):361–74.

79. Helmstetter FJ, Fanselow MSJAL, Behavior. Aversively motivated changes in meal patterns of rats in a closed economy: The effects of shock density. 1993;21(2):168–75.

80. Lesuis SL, Catsburg LAE, Lucassen PJ, Krugers HJ. Effects of corticosterone on mild auditory fear conditioning and extinction; role of sex and training paradigm. Learning & memory (Cold Spring Harbor, NY). 2018;25(10):544–9.

81. Korte SM. Corticosteroids in relation to fear, anxiety and psychopathology. Neuroscience & Biobehavioral Reviews. 2001;25(2):117–42.

